# Identification, annotation and visualisation of extreme changes in splicing from RNA-seq experiments with SwitchSeq

**DOI:** 10.1101/005967

**Authors:** Mar Gonzàlez-Porta, Alvis Brazma

## Abstract

In the past years, RNA sequencing has become the method of choice for the study of transcriptome composition. When working with this type of data, several tools exist to quantify differences in splicing across conditions and to address the significance of those changes. However, the number of genes predicted to undergo differential splicing is often high, and further interpretation of the results becomes a challenging task. Here we present SwitchSeq, a novel set of tools designed to help the users in the interpretation of differential splicing events that affect protein coding genes. More specifically, we provide a framework to identify switch events, *i.e.,* cases where, for a given gene, the identity of the most abundant transcript changes across conditions. The identified events are then annotated by incorporating information from several public databases and third-party tools, and are further visualised in an intuitive manner with the independent R package tviz. All the results are displayed in a self-contained HTML document, and are also stored in txt and json format to facilitate the integration with any further downstream analysis tools. Such analysis approach can be used complementarily to Gene Ontology and pathway enrichment analysis, and can also serve as an aid in the validation of predicted changes in mRNA and protein abundance.

Availability: The latest version of SwitchSeq, including installation instructions and use cases, can be found at https://github.com/mgonzalezporta/SwitchSeq. Additionally, the plot capabilities are provided as an independent R package at https://github.com/mgonzalezporta/tviz.

## 1 Introduction

In the past years, RNA sequencing has become the method of choice for the study of transcriptome composition. Among the strengths of this technology there is the possibility to perform a broad range of analyses without the need for intricate experimental designs: *e.g.,* identification of novel transcribed regions, deconvolution of allele specific expression and study of alternative splicing, to mention a few. Regarding this last application, several tools exist to estimate transcript expression levels (e.g. MISO (Katz *et al*., 2010), Cufflinks (Trapnell *et al*., 2010), MMSEQ (Turro *et al.*, 2011)) and to predict differences in splicing across conditions and assess their significance (e.g. MMDIFF (Turro *et al.*, 2013), Cuffdiff (Trapnell *et al.*, 2013), DEXSeq (Anders *et al.*, 2012)). Quite often though, these tools result in a long list of features (*e.g.,* genes) that is difficult to interpret. On the other hand, we recently found that gene expression is often dominated by one single transcript in protein coding genes (Gonzàlez-Porta *et al.*, 2013), and introduced the concept of *switch event* to refer to those cases where, for a given gene, the identity of the most abundant transcript changes across conditions. With both of these situations in mind, we have developed SwitchSeq, the first tool for the identification, annotation and visualisation of the most extreme and prevalent changes in splicing that affect protein coding genes.

## 2 Methods

### 2.1 Identification and annotation of switch events in protein coding genes with SwitchSeq

SwitchSeq relies on transcript level quantifications in order to identify switch events. This step can be divided into two subtasks: (i) identification of the most abundant transcript within each gene, and (ii) discovery of cases where the identity of such transcript changes across conditions. SwitchSeq takes as input a matrix of normalised counts (TPMs, FPKMs/RPKMs, etc.), thus being entirely detached from any specific mapping and quantification strategy (Figure 1). The input table can contain any number of genes and transcripts, but a useful approach is to focus on the genes for which the user has already predicted differences in splicing through other tools (e.g. MMDIFF, Cuffdiff, DEXSeq). In this way, it is possible to obtain a subset of that list that contains the most extreme and prevalent changes in splicing across the studied conditions. In addition, tools like MMDIFF provide a significance threshold for each of the tested transcripts, and such list of significant events can be passed to SwitchSeq to filter transcripts accordingly.

**Figure 1.**
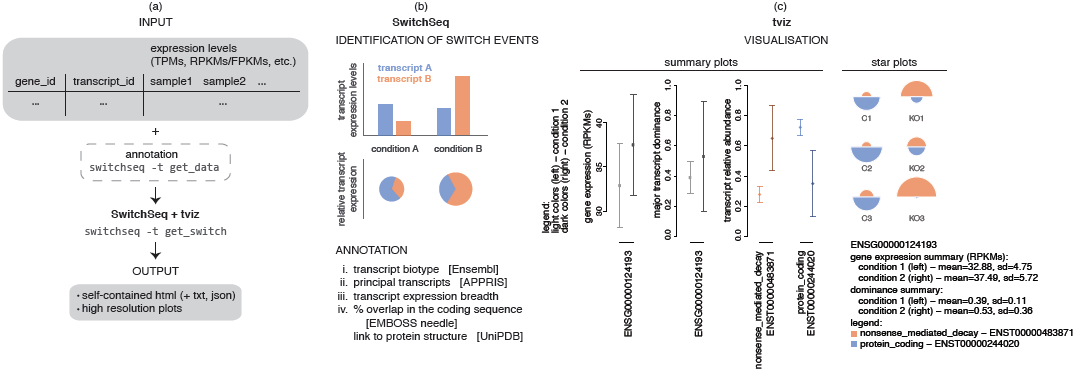
Overview of SwitchSeq. (a) The analysis workflow. SwitchSeq input consists of a matrix of normalised transcript-level counts, as well as several annotation files that can be easily obtained with the provided wrapper tool. This information is used to identify and annotate switch events, and the results are stored in a self-contained HTML report, as well as in txt and json format. Furthermore, SwitchSeq produces plots for the visualisation of the identified events, which can be easily accessed from the report. Such ploting capabilities have been implemented in an independent R package (tviz), and hence can be used independently of any SwitchSeq execution. (b) Identification and annotation of switch events. Based on the expression data provided by the user, SwitchSeq identifies the most abundant transcript within each gene and detects cases where its identity changes across conditions, here referred to as switch events. Following the identification stage, information from several public databases is incorporated to aid in the interpretation of the events. (c) Visualisation of switch events. Two types of plots are automatically generated by SwitchSeq: summary plots and star plots. Summary plots contain information on the distribution of gene expression levels, major transcript dominance and transcript relative abundances across all the samples that belong to each condition. Star plots provide information on the transcript expresison estimates in a sample-specific manner. Example plots that show a switch event between the two annotated transcripts of the human gene *SRSF6* (ENSG00000124193) are included here: the protein coding transcript ENST00000244020 is the major isoform in all the three samples of the first condition (control - C), whilst the nonsense-mediated-decay transcript ENST00000483871 becomes the most abundant isoform in the second one (knock-down - KD).

SwitchSeq combines several sources of information to facilitate the interpretation of the identified switch events, including data from public databases and results from the execution of third-party tools. This functionality depends on several annotation files, which can be easily generated by the user with the provided wrapper script (Figure 1). Specifically, the output of the annotation stage contains the following information: (i) the biotype of the transcripts involved in the switch, which is retrieved from Ensembl (Flicek *et al.*, 2013) and can be informative of their function (e.g. the most common biotypes for transcripts derived from protein coding genes are *protein coding*, *nonsense mediated decay*, *retained intron* and *processed transcript*); (ii) whether the identified transcripts are classified as principal isoforms in APPRIS (Rodriguez *et al.*, 2013), a resource that aims at selecting a set of representative transcripts for each protein coding gene based on conservation data and protein structural and functional information; (iii) the expression breadth for each of the identified transcripts, i.e. the number of samples in which the transcript is detected as major relative to the number of samples in which the gene is expressed (calculated by SwitchSeq); and (iv) in the cases in which the switch involves two protein coding transcripts, the overlap in their coding sequences (calculated by SwitchSeq by performing a global alignment with EMBOSS Needle (Rice *et al.*, 2000)), as well as the protein sequences themselves for further analysis of potential functional differences with tools like MAISTAS (Floris *et al.*, 2011), and links to the available structures for the protein, if any, including a graphic representation of their coverage (through the UniPDB widget).

### 2.2 Visualisation of switch events with tviz

For each switch event, two types of plots are generated to enable visual inspection of the predicted changes (Figure 1). On the one hand, summary plots include combined information on the expression levels across all the samples in each condition. More specifically, they include information on the distribution of gene expression levels, relative abundances for all the annotated transcripts and major transcript dominance (*i.e.,* difference in the abundance of the first *vs.* the second most abundant transcripts of each gene). These data are represented in the form of box plots or as the average plus error bars depending on the number of samples available. On the other hand, sample-specific visualisation of transcript expression levels is achieved with star plots. In such plots each sample is plotted as a pie chart, where transcript abundance is represented by the size of each slice, and gene expression is represented by the overall size of the plot, thus allowing comparisons across samples. Both types of plots can be produced for any gene and independently of SwitchSeq execution with the R package tviz.

### 2.3 SwitchSeq output

The identified switch events and associated information are reported in a self-contained HTML document, where the results can be easily filtered, formatted and sorted (including ranking to maximise expression breadth in both conditions). Such HTML also contains additional information on the options used during the execution. Finally, the results are also available in txt and json format, to facilitate the integration with any further downstream analysis tools.

## 3 Conclusion

Evaluating differences in splicing across conditions has become a common task in any RNA-seq data analysis pipeline. However, the number of genes predicted to undergo changes in splicing is often high, and further interpretation of the results becomes a challenging task. SwitchSeq has been designed to help the users in the interpretation of differential splicing events that affect protein coding genes, by letting them focus on the most extreme and prevalent changes. The identified switch events are further annotated through the combination of expression data and information retrieved from several public databases, and can be also visualised in an intuitive manner. Such analysis approach can be used complementarily to Gene Ontology and pathway enrichment analysis, can serve as an aid in the validation of predicted changes in mRNA or protein abundance, and can also be applied to investigate the functional impact of the detected alterations in splicing across conditions (*e.g.,* healthy *vs.* tumour samples). SwitchSeq has been written in Perl and R and is available as open source software, thus enabling further customisation.

## Acknowledgements

We would like to aknowledge the members of the Functional Genomics team for their feedback and suggestions, specially Nuno Fonseca, Natalja Kurbatova, Mitra Barzine, Liliana Greguer and Jing Su, as well as Ernest Turró.

